# MuSK antibodies differently affect the MuSK signaling cascade depending on valency and epitope specificity

**DOI:** 10.64898/2026.03.17.709302

**Authors:** Dana L.E. Vergoossen, Robyn Verpalen, Stine Marie Jensen, Sandra Fonhof, Yvonne E. Fillié-Grijpma, Christoph Gstöttner, Elena Dominguez-Vega, Silvère M. van der Maarel, Jan J.G.M. Verschuuren, Maartje G. Huijbers

## Abstract

Muscle-specific kinase (MuSK) is a pivotal player in forming and maintaining healthy neuromuscular junctions (NMJ). In MuSK myasthenia gravis (MG), autoantibodies targeting MuSK disrupt its function, impairing neuromuscular transmission and causing fatigable skeletal muscle weakness. MuSK autoantibodies predominantly belong to the IgG4 subclass, which bind in a monovalent fashion due to Fab-arm exchange, although autoantibodies of other subclasses also exist. Polyclonal autoreactive IgG from patients may therefore harbor a variety of monovalent and bivalent MuSK antibodies with potentially distinct effects on MuSK signaling. To further unravel the pathomechanisms underlying MuSK MG, we have investigated how MuSK antibody-binding affects MuSK functioning with a diverse panel of (patient-derived) monoclonal MuSK antibodies. Our findings reveal that the valency of antibody-binding influences binding kinetics to MuSK, inhibition of agrin-induced MuSK activation, Dok7 binding to MuSK and NMJ gene expression. Monovalent binding to the frizzled domain of MuSK did not inhibit agrin-induced MuSK activation, while monovalent binding to the Ig-like domain 1 does. Moreover, the kinetics of Dok7 degradation induced by bivalent MuSK antibodies appear to depend on binding-epitope of MuSK. Surprisingly, none of the clones tested (both bivalent and monovalent) increased MuSK internalization. Taken together, the cumulative pathogenic effect of polyclonal MuSK antibodies in individual MuSK MG patients thus likely depends on autoantibody titer, affinity and the unique composition of MuSK autoantibodies varying in epitope and valency. This research enriches our understanding of the intricate interactions between antibodies and MuSK in MuSK MG and offers potential insights into novel therapeutic strategies using MuSK antibodies.

## 1. Introduction

The neuromuscular junction (NMJ) is a specialized synapse where motor neurons and skeletal muscles communicate. Many proteins in the NMJ are indispensable as neuromuscular communication is essential for basic muscular functions such as breathing [1]. Muscle-specific kinase (MuSK) is a transmembrane tyrosine kinase which forms a signaling hub essential for forming and actively maintaining the NMJ [1]. MuSK has three extracellular immunoglobulin-like (Ig-like) domains followed by a frizzled-like domain (Fz-domain), a transmembrane domain and an intracellular kinase domain [2]. The interaction between the MuSK Ig-like domain 1 and lipoprotein receptor-related protein 4 (Lrp4) is crucial for activation and autophosphorylation of the MuSK kinase domain through dimerization of two MuSK molecules [3, 4]. The interaction between Lrp4 and MuSK is greatly enhanced once neuronal agrin released from the motor nerve terminal has bound Lrp4 [4]. Upon MuSK dimerization and autophosphorylation, downstream of kinase 7 (Dok7) is recruited to the intracellular phosphotyrosine-binding site, stabilizing and enhancing phosphorylation of both MuSK and Dok7 [5, 6]. This initiates further downstream signaling leading to cytoskeletal reorganization and clustering of acetylcholine receptors (AChRs) by scaffold protein rapsyn, and contributes to NMJ-specific gene regulation [1, 7]. In addition, Collagen Q (ColQ) is involved in tethering acetylcholine esterase (AChE) to the NMJ through Lrp4 and may regulate MuSK activation by competing with agrin [8]. AChE breaks down acetylcholine, the main neurotransmitter responsible for neurotransmission at the NMJ and thereby regulates neuromuscular communication. Because MuSK has multiple functions in the NMJ, perturbing or modifying its functioning can have many consequences.

Antibodies against MuSK cause the neuromuscular autoimmune disorder myasthenia gravis (MuSK MG), characterized by fatigable skeletal muscle weakness [9, 10]. MuSK autoantibodies are predominantly of the IgG4 subclass and thus become bispecific and functionally monovalent through the stochastic process of Fab-arm exchange in the circulation [11-14]. Although the great majority of IgG4 is estimated to be bispecific, varying amounts of bivalent IgG4 MuSK antibodies have been detected in patients [12, 15, 16]. Besides IgG4 MuSK antibodies, some patients also have low levels of co-occurring functionally bivalent and monospecific IgG1, IgG2 or IgG3 MuSK antibodies [11, 14, 17]. For clarity throughout this study, the term “bivalent” will be used for functionally bivalent and monospecific antibodies, and the term “monovalent” for functionally monovalent and bispecific antibodies.

Bivalent MuSK antibodies can (partially) activate MuSK (agonists), while monovalent MuSK antibodies inhibit agrin-induced MuSK activation (antagonists) [18-20]. These opposing effects are related to the natural dimerization of MuSK that occurs in a healthy synapse to keep the MuSK kinase constitutively active and the NMJ intact. IgG4 MuSK antibodies block Lrp4-MuSK interaction thereby preventing dimerization and activation of the kinase and AChR clustering leading ultimately to synaptic disintegration and skeletal muscle fatigue [17, 21]. In contrast, bivalent MuSK antibodies are thought to force dimerization of MuSK, therefore bypassing the need of agrin and Lrp4 in activating MuSK [18, 20, 22, 23]. Experiments with recombinant monoclonal antibodies based on anti-MuSK B-cell receptor (BCR) sequences isolated from MuSK MG patients show that monovalent MuSK antibodies are more pathogenic than their bivalent equivalents and cause rapid onset fatigable muscle weakness in mice [19]. However, the pathogenicity of bivalent MuSK antibodies varies between clones. Although their AChR clustering capabilities are similar *in vitro*, some bivalent MuSK antibodies can cause myasthenic muscle weakness in mice (with slower disease progression compared to monovalent MuSK antibodies), while others remain non-pathogenic even after long-term exposure [18, 19, 24, 25]. We therefore hypothesize that the mechanism by which MuSK antibodies impair MuSK signaling differs between clones and may depend on antibody valency and epitope specificity.

Here we investigated the effect of six MuSK antibody clones in monovalent and bivalent format on MuSK binding kinetics, Dok7 binding and Dok7 levels, MuSK internalization, and NMJ gene expression.

## 2. Material and methods

### 2.1. mAb production and cFAE

Anti-MuSK clones binding to the Ig-like domain 1 of MuSK were previously isolated from a MuSK MG patient [18]. Together with mAb13 as a MuSK Fz domain binder, these clones were produced in an IgG4 Fc tail with the S228P amino-acid change and their original light chain (GeneArt, [19, 26, 27]). The b12 antibody suitable for cFAE was used as an exchange partner and control antibody [19]. Recombinant bivalent and monovalent monoclonal antibodies (mAbs) were produced, quantified and assessed on quality as described previously [18, 19]. Recombinant antibodies 11-3F6, 13-3B5 and b12 were produced in CHO cells. The other MuSK mAbs were produced in HEK cells.

### 2.2. Production of recombinant hMuSK-TST

A plasmid containing hMuSK extracellular domain (aa 1 – 486, NCBI protein: EAW59066.1), an N-terminal sortase motif, G4S linker and TwinStrepTag (SA-WSHPQFEK-(GGGS)2-GGSA-WSHPQFEK) (hMuSK-TST) was ordered in the pFastBac vector (Twist). This plasmid was transformed into a bacmid using DH10Bac competent *E. coli.* Colonies were grown for 3 days on agar plates containing gentamycin (7 µg/mL), IPTG (40 µg/mL), kanamycin (50 µg/mL), tetracyclin (10 µg/mL) and X-gal (80 µg/mL) before blue/white screening. Bacmid transposition (yielding a 3.9 kb band) was confirmed by DreamTAQ PCR using M13 primers on minipreps. The bacmid was transfected into Sf9 (*Spodoptera frugiperda)* cells (0.5E6 cells in 6-well) using 10 μl bacmid solution, 7 μl CellFectin (10362100, ThermoFisher) in 200 μl SFM-II medium (10902096, ThermoFisher) according to manufacturer’s instructions for 5 hours at 27°C before replacing medium. Supernatant containing virus was harvested after 48 hours and used 2 mL supernatant to infect 50 mL of Sf9 cells (1E6 cells/mL). Supernatant was harvested after 72 hours and stored at 4°C.

For hMuSK-TST protein production, 500 mL containing 5 E8 Tni (*Trichoplusia ni*) insect cells were transduced with 3 mL virus-containing supernatant and incubated at 27°C for 3-4 days. Supernatant cleared from cells using centrifugation (5000 g), supplemented with 10% v/v 1M Tris HCl pH 8, 2.5% v/v 5M NaCl, 0.2% v/v 0.5M EDTA, 0.5% BioLock (2-0205-050, IBA), the pH adjusted to 8 and filtered (0.22 μm). Affinity purification was done on an AKTA pure (Cytiva) using a Strep-Tactin®XT 4Flow® high capacity FPLC column (2-5027-001, IBA) according to manufacturer’s instructions. Briefly, the column was equilibrated with wash buffer (100mM Tris HCl, pH 8, 150 mM NaCl, 1mM EDTA) for 10 column volumes (CV) before loading the sample at 1 mL/min. After loading the column was washed with wash buffer for 5 CV and hMuSK-TST was eluted with elution buffer (50 mM biotin in wash buffer). Protein-containing fractions were dialysed to PBS, sterile filtered, flash frozen in liquid N_2_ and stored at -80°C.

### 2.3. SPR

All SPR experiments were performed using a Biacore T200, with a 1 Hz data collection rate at 25 °C, using 10 mM HEPES, 150 mM NaCl, 3 mM EDTA, 0.005% P20, pH 7.4 as running buffer. To assess the affinity and binding kinetics of MuSK antibodies to MuSK, a protein A sensor chip (29127555, lot10308130, Cytiva) was used to capture the purified recombinant bivalent antibodies (0.5 nM in running buffer, ∼150 kDa) to a density of ∼ 28RU on flow cell 2 at 10 µL/min and flow cell 1 was left blank to serve as a reference surface. To collect kinetics data, hMuSK-TST in running buffer, was injected over the two flow cells at 0.3, 0.9, 2.7, 8.1 and 24.3 nM at a flow rate of 30 µL/min using single-cycle kinetics with an association time of 120 s per concentration and a dissociation time of 1700 s. The surfaces were regenerated with 20 s of 10 mM Glycine HCl, pH 1.5 at 20 µL/min. Sample and blank cycles were run in duplicate per clone within each experiment and at least three independent experiments were done. Per independent experiment, cycles with artifacts were removed and the data were fit to a 1:1 interaction model using the global analysis option available within the Biacore T200 Evaluation Software (Cytiva, version 3.2.1).

To assess the binding of both monovalent and bivalent MuSK antibodies to MuSK, a CM5 sensor chip (29104988, lot10328745, Cytiva) was immobilized with StreptactinXT (13 kDa) using amine coupling according to the kit instructions (2-4370-000, Cytiva) to a density of 1501 RU on flow cell 3 and 1646 RU on flow cell 4. To collect binding data hMuSK-TST (50 ng/mL in 10 mM acetate pH 4.5) was captured on flow cell 4 for 80 s at 10 µL/min to reach a density of ∼ 18 RU with a stabilization period of 60 s. The recombinant antibodies (∼ 150 kDa) were injected over both flow-cells at 0.33, 1, 3, 9 nM in running buffer using single-cycle kinetics at 30 µL/min using a 300 s association time and 1700 sec dissociation time. The surface was regenerated with three pulses of 60 s of 3 M GuHCl at 10 µL/min with 30 sec stabilization and an extra running buffer wash. Sample cycles were run in duplicate within each experiment with duplicate blank cycles every two sample cycles. Cycles with artifacts were removed and the dissociation phases of each sample were fit with a 1:1 interaction model using the the global analysis option available within the Biacore T200 Evaluation Software (Cytiva, version 3.2.1). Three independent experiments were done.

### 2.4. C2C12 culturing and treatment conditions

C2C12 myoblasts were obtained from CLS Cell Lines Service GmbH (Eppelheim, Germany), tested for mycoplasma contamination and maintained for maximum 5 passages after thawing. Myoblasts were grown in proliferation medium (DMEM Glutamax (10566016, Thermo Fisher) supplemented with 10% FBS (S1810-500, Biowest) and 1% penicillin/streptomycin (15140122, Gibco). Cells were plated at 1.25e^4^ – 2e^4^ cells per cm^2^ in proliferation medium. Once cells reached 90 – 95 % confluency, differentiation was induced by DMEM Glutamax supplemented with 2% HS and 1% penicillin/streptomycin, refreshed every 2-3 days. Experiments were done on day 5 of differentiation. All mAbs were used at 7.7nM, neural agrin (550-AG-100, R&D systems) at 0.1nM, neuregulin (396-HB-050, R&D systems) at 4.9nM, EGF at 200ng/mL (236-EG-200, R&D systems), unless otherwise specified.

### 2.5. MuSK immunoprecipitation

MuSK was immunoprecipitated (IP) as described previously [19]. Briefly, differentiated C2C12 myotubes were cultured in 10cm dishes, treated for 30 min and lysed in phosphate lysis buffer (30 mM triethanolamine, 1% NP 40, 50mM NaF, 2mM sodium orthovanadate, 1mM sodium tetrathionate, 5 mM EDTA, 5 mM EGTA, 1mM N-ethylmaleimide, 50mM NaCl, 1× protease inhibitor cocktail, 1× phosphatase inhibitor cocktail). Lysates were cleared by centrifuging for 20 min at 5.000 g, adjusted to ensure equal protein input over all conditions and combined with 1µg/sample 11-3F6 IgG1 to IP MuSK. Antibody-protein complexes were captured with protein A agarose beads (11134515001, Roche) and eluted with 40-50μL 2x sample buffer (40mM Tris-HCl pH6.8, 3.3% SDS, 16.5% glycerol, 0.005% Bromophenol blue, 0.2M DTT) and incubated at 95 degrees for 5 min.

### 2.6. Protein isolation

To assess Dok7 levels, C2C12 myotubes were cultured in 6-well plates. After treatment, myotubes were harvested in ice-cold PBS and stored as cell pellets at -80 prior to protein isolation. Pellets were lysed in RIPA buffer (50mM Tris pH 7.4, 150mM NaCl, 1mM EDTA, 0.1% SDS, 1% NP-40, 0.5% sodium deoxycholate, 1x protease inhibitor cocktail) and cleared by centrifuging for 20 min at 5.000 g. Protein content of lysate was determined using BCA protein assay kit (23225, Thermo Scientific) and used to prepare samples with equal protein content in sample buffer.

### 2.7. Surface depletion assay

Immediately after treatment exposure, cells were put and kept on ice in the cold room (4°C) until lysis. Cells were thoroughly washed with ice cold PBS^2+^ (1.5mM MgCl_2_; 0.2mM CaCl_2_ in PBS, pH 7.4). Freshly prepared 1mg/mL Sulfo-NHS-SS-Biotin (PG82077, Thermo Fisher) in PBS^2+^ was added to each plate and incubated shaking vigorously (∼300 rpm) for 30 min. Unbound Sulfo-NHS-SS-Biotin was washed away with quenching buffer (100mM Glycine in PBS^2+^) by rinsing three times and incubating shaking for 2x 15 minutes. All cells were washed three times with ice cold PBS^2+^before being lysed with RIPA buffer (10 mM Tris, pH 7.4; 150 mM NaCl; 1 mM EDTA; 0.1% SDS; 1.0% triton X-100; 1.0% sodium deoxycholate, 1x protease inhibitor cocktail). Lysates of all samples were adjusted to ensure equal protein input over all conditions based on BCA quantification. Adjusted lysates were incubated with streptavidin beads (20349, Thermo Scientific) rotating ON (4°C). Protein was eluted from the beads using 2x sample buffer.

### 2.8. Western blotting

Protein samples were ran on SDS-PAGE gel and transferred to PDVF membrane. Western blot conditions for different sample types can be found in Appendix A, Table S1. Chemiluminescence was measured on the Amersham Imager 600 (Cytiva). Immunofluorescence was measured with the Odyssey (Licor).

### 2.9. RNA isolation

C2C12 myotubes or frozen muscle tissue were lysed and homogenized in QIAzol lysis reagent (Qiagen). Total RNA was extracted and purified with the miRNeasy Mini Kit according to manufacturer’s instructions (Qiagen, 1038703). RNA was treated with DNase (Qiagen) on column for 30 min at room temperature. RNA concentration was quantified using NanoDrop ND-1000 spectrophotometer (Thermo Fisher Scientific).

### 2.10. Quantitative Real-time PCR

First strand cDNA was synthesized from 1000-3000 ng total RNA with the RevertAid H Minus First Strand cDNA Synthesis kit using oligo(dT) primers (Thermo Fisher Scientific, K1632). Relative gene expression levels were determined with iQ SYBR Green Supermix (Bio-Rad, #1708886) and 1 pM forward and reverse primers (Appendix A, Table S2) on CFX384 Touch Real-Time PCR Detection System (Bio-Rad) with the following program: 95°C for 3 min, 40 cycles of 10 s at 95°C and a melting temperature of 60°C for 30 s, followed by a melting curve analysis from 65°C to 95°C (temperature increments of 0.5°C). Quantification cycle (Cq) values were obtained from CFx manager or maestro software (BioRad). All samples were run in triplicate and *Gapdh* and *Rpl13a* were used as housekeeping genes. Technical replicates that differed >0,5 in Cq from the others in the triplicate were excluded. Normalized fold changes were calculated compared to untreated or b12 using the validated efficiency of each primer (Appendix A, Table S2).

### 2.11. Bulk RNA sequencing and analysis

Total RNA integrity of the 16 h-treated C2C12 myotube samples was analyzed with the Agilent BioAnalyzer RNA Nano 6000 chip and all had an RNA Integrity Number of > 9.5 (Agilent Technologies, Amstelveen, the Netherlands). The library was prepared with the TruSeq Stranded Total RNA with Ribo-Zero H/M/R kit and 30 million reads were sequenced with the NovaSeq 6000 PE 150 system (Illumina) by Macrogen. Reads were trimmed and quality filtered by TrimGalore (v.0.6.6, Cutadapt v.2.10), using default parameters to remove low-quality nucleotides. Consequently, reads were mapped to the Genome Reference Consortium Mouse Build 38, using STAR Aligner (v.2.7.6a). A gene expression counts table was generated with HTSeq (v.0.12.4). Data were normalized for sequencing depth with the median of ratios method and consequently analyzed for differential expression in the DESeq2 R package (v.1.32.0). Genes with an adjusted p-value < 0.05 (Benjamini-Hochberg) were considered significant. The RNA-seq data have been deposited in NCBI’s Gene Expression Omnibus and are accessible through GEO Series accession number GSE 316686 (https://www.ncbi.nlm.nih.gov/geo/query/acc.cgi?acc=GSE 316686) [28].

### 2.12. Statistics

Statistical analyses were done in GraphPad Prism software (version 9.3.1) or R (version 4.3.1). Which statistical test was used per experiment is described in the figure legends. Data is presented as (geometric) mean with (geometric) standard error of the mean. p values <0.05 were considered significant. ∗p < 0.05; ∗∗p < 0.01; ∗∗∗p < 0.001; ∗∗∗∗p < 0.0001.

## 3. Results

### 3.1. Generation and quality control of bivalent and functionally monovalent antibodies

To investigate how MuSK antibodies with different characteristics affect MuSK-mediated signaling, we generated a panel of six human recombinant MuSK antibodies (Table 1). The sequences of five MuSK clones were isolated from MuSK MG patient BCR sequences from one patient and mAb13 was previously identified by others after phage display selection and screening [18, 26, 27]. All clones were produced in a bivalent manner. Monovalent equivalents were generated by IgG4 controlled Fab-arm exchange with the b12 control antibody and are described as [clone]xb12 [19]. The residual amount of bivalent MuSK antibody after the controlled Fab-arm exchange method was two percent or less, validating the relative purity of the monovalent MuSK antibodies (Appendix A, Figure S1A). The method and purity were previously validated for 11-3F6xb12 and 13-3B5xb12 [19, 29]. Due to Fab-glycosylation, the 11-3D9xb12 mixture was too complex to calculate relative amounts of each variant. However, the deconvoluted mass spectra in this mixture confirmed the generation of the bispecific, monovalent variant for this clone (Appendix A, Figure S1B). Five out of six clones bind the Ig-like domain 1 of MuSK, while one clone (mAb13) binds the Fz domain [18, 30]. Within the Ig-like domain 1, the 13-3B5 clone binds a non-overlapping epitope compared to the other Ig-like domain 1 binding clones [24].

**Table 1:** Panel of anti-MuSK clones with antibody characteristics.

| MuSK clone | MuSK domain | $K_D$ (M)<sup>*</sup> | $k_a$ (M<sup>-1</sup>s<sup>-1</sup>)<sup>*</sup> | $k_d$ (s<sup>-1</sup>)<sup>*</sup> | Functional Valency | *In vivo* muscle weakness |
| --- | --- | --- | --- | --- | --- | --- |
| 13-3B5 | Ig-like 1 | 1.4 10<sup>-12</sup> | 1.6 10<sup>6</sup> | 2.2 10<sup>-6</sup> | Bi | Slow<sup>1</sup> |
|  |  |  |  |  | Mono | Rapid<sup>1</sup> |
| 11-3F6 | Ig-like 1 | 4.4 10<sup>-12</sup> | 2.7 10<sup>6</sup> | 1.2 10<sup>-5</sup> | Bi | No<sup>1</sup> |
|  |  |  |  |  | Mono | Rapid<sup>1</sup> |
| 13-3D10 | Ig-like 1 | 3.8 10<sup>-11</sup> | 3.8 10<sup>6</sup> | 1.5 10<sup>-4</sup> | Bi | No<sup>2</sup> |
|  |  |  |  |  | Mono | n.a. |
| 11-3D9 | Ig-like 1 | 2.5 10<sup>-11</sup> | 1.7 10<sup>6</sup> | 4.4 10<sup>-5</sup> | Bi | No<sup>2</sup> |
|  |  |  |  |  | Mono | n.a. |
| 13-4D3 | Ig-like 1 | 7.4 10<sup>-11</sup> | 1.9 10<sup>6</sup> | 1.4 10<sup>-4</sup> | Bi | No<sup>2</sup> |
|  |  |  |  |  | Mono | n.a. |
| mAb13 | Fz | n.a. | n.a. | n.a. | Bi | No<sup>2</sup> |
|  |  |  |  |  | Mono | n.a. |

<sup>1</sup>Vergoossen et al. 2021 PNAS
<sup>2</sup>Lim et al. 2023 Sci Rep.
\*Geometric mean of n=3-4, measured using SPR on human MuSK
$k_a$ = association rate constant, $k_d$ = dissociation rate constant, $K_D$ = equilibrium dissociation constant, SPR = surface plasmon resonance

### 3.2. MuSK antibodies vary in affinity and binding kinetics depending on clone and valency

The kinetics of MuSK antibody binding was assessed with surface plasmon resonance (SPR) by capturing the antibodies on the chip with protein A and using the extracellular domain of human MuSK tagged with TwinStrepTag (hMuSK-TST) as analyte (Appendix A, Figure S2A). The clones binding the Ig-like domain 1 bind human MuSK with equilibrium dissociation constants (*K_D_*) in the picomolar range, indicating they have very high affinity (Table 1, Appendix A, Figure S2E). The variation in affinity between the clones is mainly driven by differences in the dissociation rate constant, *k_d_* (Table 1, Appendix A, Figure S2D), varying by two orders of magnitude. The *k_d_* for 13-3B5 was below the limit of detection for the Biacore T200 despite the extended dissociation time of 1700 s, indicating 13-3B5 does not detectably release from MuSK once it is bound. This means both the *k_d_* and *K_D_* should be interpreted as estimations of very low constants for the 13-3B5 clone. The association rate constants (*k_a_*) of the different clones vary less and lie within a two-fold difference of each other (Appendix A, Figure S2C). The binding kinetics of mAb13 were not assessed, as it was shown it does not bind to human MuSK [24].

Although capturing the antibodies on the chip is the gold standard for assessing the affinity of antibodies with the equilibration dissociation constant, it is not suitable to assess the effects of antibody valency on the binding kinetics. In addition, MuSK is immobilized in the membrane in muscle, which is not reflected in the protein A capture assay. To better model the effects of MuSK autoantibodies binding to synaptic MuSK, we captured hMuSK-TST on the chip using immobilized StreptactinXT and used the monovalent and bivalent variants of the MuSK antibodies as analytes (Figure 1A). Because bivalent antibody binding gives avidity effects, only the dissociation phases are fitted with a 1:1 binding model to quantify the apparent dissociation rate constant (Figure 1B and C). The apparent *k_d_* of the monovalent variants was significantly higher than the bivalent variants (F(1,20) = 561.14, p < 0.0001), confirming that monovalent MuSK antibodies are more easily released from MuSK than their bivalent equivalents. The apparent *k_d_* also significantly depended on the clone (F(4,20) = 47.7, p < 0.0001), with 13-3B5 having significantly slower dissociation than all other clones and 11-3F6 having faster dissociation than 13-3B5, but slower dissociation than 11-3D9, 13-3D10 and 13-4D3 (Appendix A, Table S3). Taken together, all clones show minimal dissociation in bivalent form with 13-3B5 having the slowest dissociation in both monovalent and bivalent form, supporting their very high affinity for MuSK as measured above.

**Figure 1.**
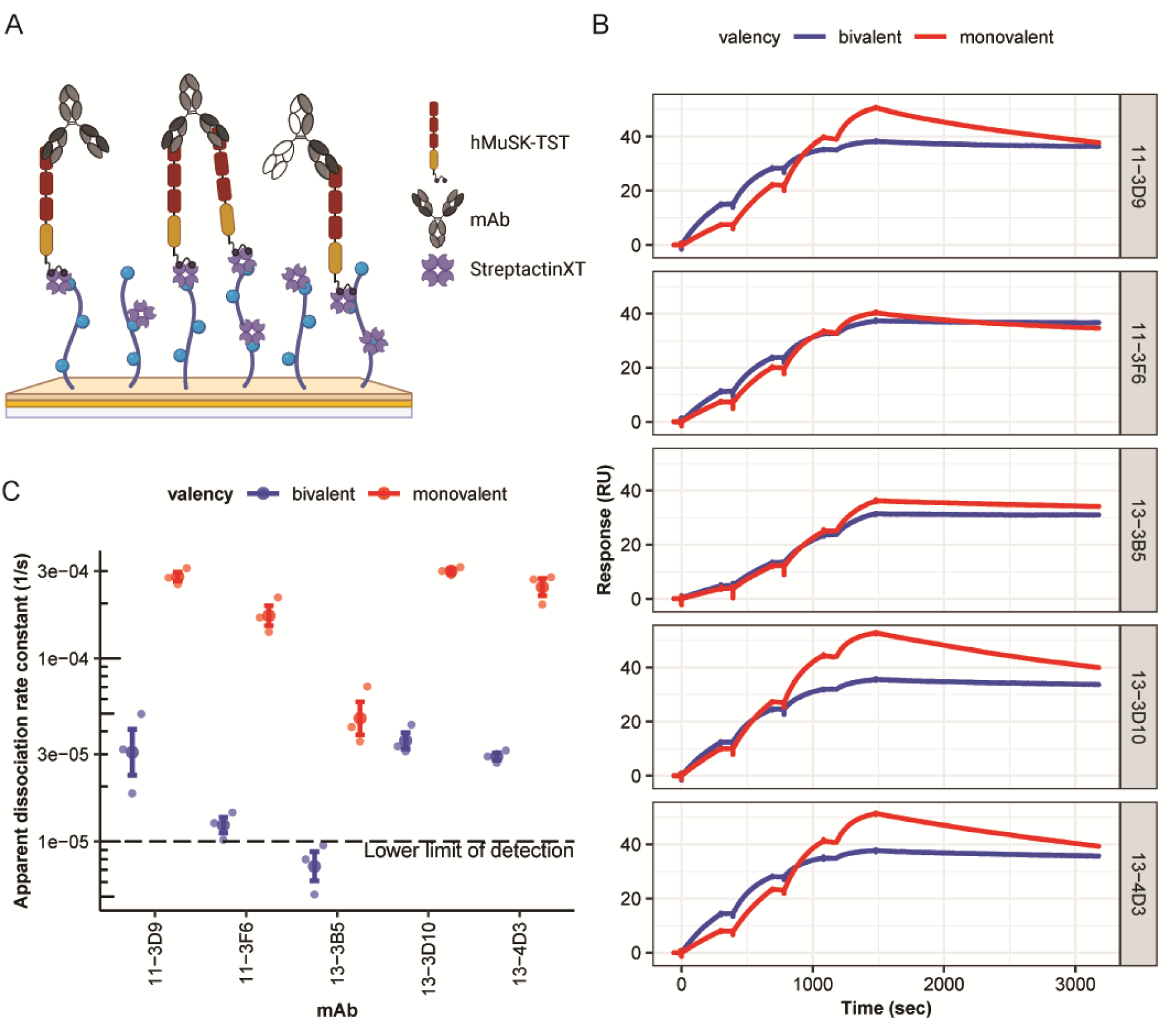
Binding kinetics of MuSK antibodies to MuSK vary depending on valency and clone. (A) Graphical representation of SPR assay with hMuSK-TST as ligand, captured with StreptactinXT on the chip, and bivalent or monovalent mAbs as analyte. (B) Representative double referenced sensorgrams of binding of monovalent (red) or bivalent (blue) MuSK antibodies to MuSK. (C) Dissociation-rate constants fitted with a 1:1 model to only the dissociation phase. Data represents geometric mean and geometric SEM of n=3 independent replicates. The effects of valency and clone was assessed by two-way ANOVA on the log10-transformed values.

### 3.3. MuSK antibodies differentially affect MuSK and Dok7 depending on valency and epitope

C2C12 myotube cultures are a well-established cell model to interrogate MuSK signaling *in vitro* as they endogenously express the muscle-specific signaling components of this pathway. All five monovalent MuSK antibodies binding the Ig-like domain 1 of MuSK fully inhibited agrin-induced MuSK phosphorylation, while their bivalent equivalents do not (Figure 2A and B). Monovalent mAb13xb12 binding the Fz domain of MuSK did not inhibit MuSK phosphorylation, even at higher concentrations (Figure 2A, 2B and Appendix A, S3A). Bivalent mAb13 was able to induce MuSK phosphorylation in the absence of agrin, while monovalent mAb13xb12 was not (Appendix A, Figure S3B). For agrin in combination with bivalent 13-3B5 or mAb13, MuSK phosphorylation exceeded the levels of agrin with the b12 control antibody, while for bivalent 13-3D10 the MuSK phosphorylation levels were slightly lower (Figure 2A and B). This suggests that the bivalent MuSK antibody is facilitating MuSK phosphorylation over agrin. Taken together, the inhibition of MuSK by monovalent MuSK antibodies seems to depend on which the structural domain of MuSK is targeted, while the agonistic capacity on MuSK phosphorylation of bivalent MuSK antibodies does not seem to depend on these epitopes.

**Figure 2.**
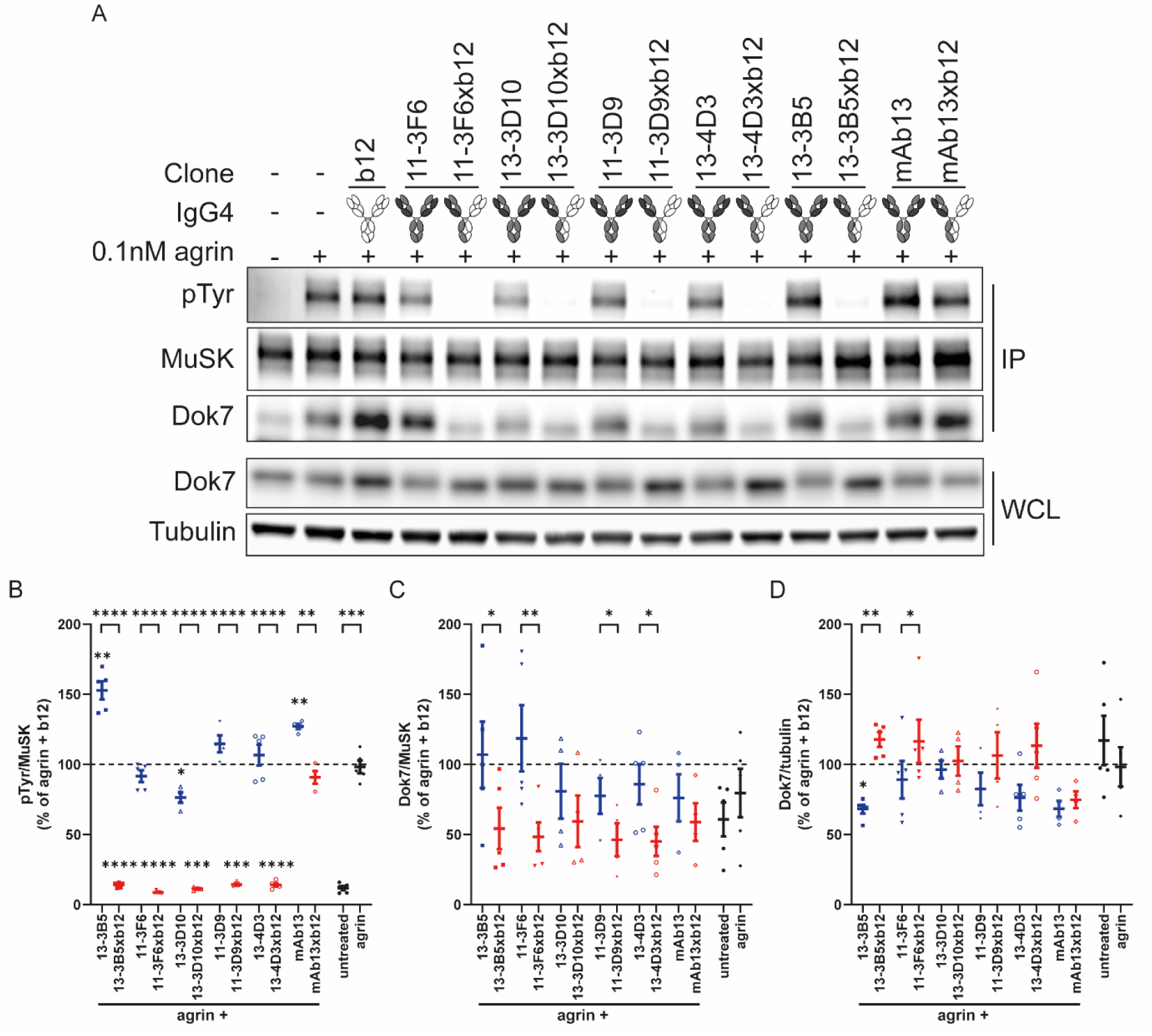
MuSK antibodies differentially affect MuSK and Dok7 depending on valency and epitope. (A) Representative blots of MuSK phosphorylation signal and MuSK-Dok7 co-immunoprecipitation. Quantification of (B) phosphorylated MuSK, (C) Dok7 interacting with MuSK and (D) Dok7 levels in whole cell lysate after 30 min exposure. Data represents mean and SEM. n=5 for 13-3B5(xb12), 11-3F6(xb12), 13-4D3(xb12) and agrin and n=4 for 11-3D9(xb12), 13-3D10(xb12) and mAb13(xb12). Paired t-test on log-transformed data with Benjamini-Hochberg false discovery rate correction. IP = immunoprecipitation, WCL = whole cell lysate. ∗p < 0.05; ∗∗p < 0.01; ∗∗∗p < 0.001; ∗∗∗∗p < 0.0001.

Dok7 binding to MuSK is critical to stabilize and enhance MuSK phosphorylation and propagate intracellular signaling [5, 6]. If Dok7 cannot properly bind to phosphorylated MuSK, this can contribute to antibody pathogenicity. To assess how much Dok7 interacts with MuSK upon MuSK antibody-binding, endogenously expressed MuSK was immunoprecipitated and assessed for co-immunoprecipitation of Dok7. Inhibition of MuSK phosphorylation by monovalent MuSK antibodies resulted in less Dok7 bound to MuSK compared to their bivalent equivalents for the MuSK Ig-like 1 binding 13-3B5, 11-3F6, 11-3D9 and 13-4D3 clones (Figure 2A and C). No large differences were found in the recruitment of Dok7 to MuSK between monovalent and bivalent variants for the 13-3D10 and mAb13 clones, or between individual bivalent or monovalent anti-MuSK clones. To investigate if equal amounts of Dok7 were available for binding to MuSK in all conditions, we investigated total Dok7 levels in whole cell lysate (WCL). Bivalent 13-3B5 and 11-3F6 significantly reduced Dok7 protein levels compared to agrin with their monovalent equivalents (Figure 2A and D). In addition, agrin with bivalent 13-3B5 has significantly lower Dok7 levels compared to agrin with control antibody b12. Tendencies for lower Dok7 levels compared to agrin were also seen for some of the other, especially bivalent, MuSK clones.

To further investigate the kinetics of how MuSK activation by bivalent MuSK antibodies or agrin influences Dok7, we measured Dok7 levels in whole cell lysate of treated C2C12 cultures over time. Bivalent mAb13 reduced Dok7 most similarly to agrin, starting at 2 hours and continuing to 6 hours (Figure 3). In contrast, bivalent MuSK clones binding the Ig-like domain 1 already significantly reduced Dok7 levels after 30 minutes, but do not appear to further decrease Dok7 levels at later time points (Figure 3). Consistently, Dok7 levels upon 30-minute exposure to bivalent 13-3B5, 13-3D10 and 13-4D3 were significantly lower than mAb13 (Figure 3C). Bivalent 13-3B5 reduced Dok7 levels more than all other Ig-like domain 1 binding clones at 2 and 6 hours and compared to bivalent 11-3F6 at 30 minutes (Figure 3C-E). Bivalent mAb13 induced significantly lower Dok7 levels compared to the 13-3D10 and 13-4D3 clones at 6 hours (Figure 3E). Agrin induced significantly lower Dok7 levels compared to bivalent 13-3D10 at 6 hours, but no other significant differences between agrin and the bivalent MuSK mAbs could be detected after multiple testing correction (Figure 3E). Since mAb13 binds the Fz domain and the 13-3B5 clone binds a different epitope compared to the other Ig-like domain 1 binding clones, Dok7 levels appear to be affected differently by bivalent MuSK antibodies depending on antibody epitope [24].

**Figure 3.**
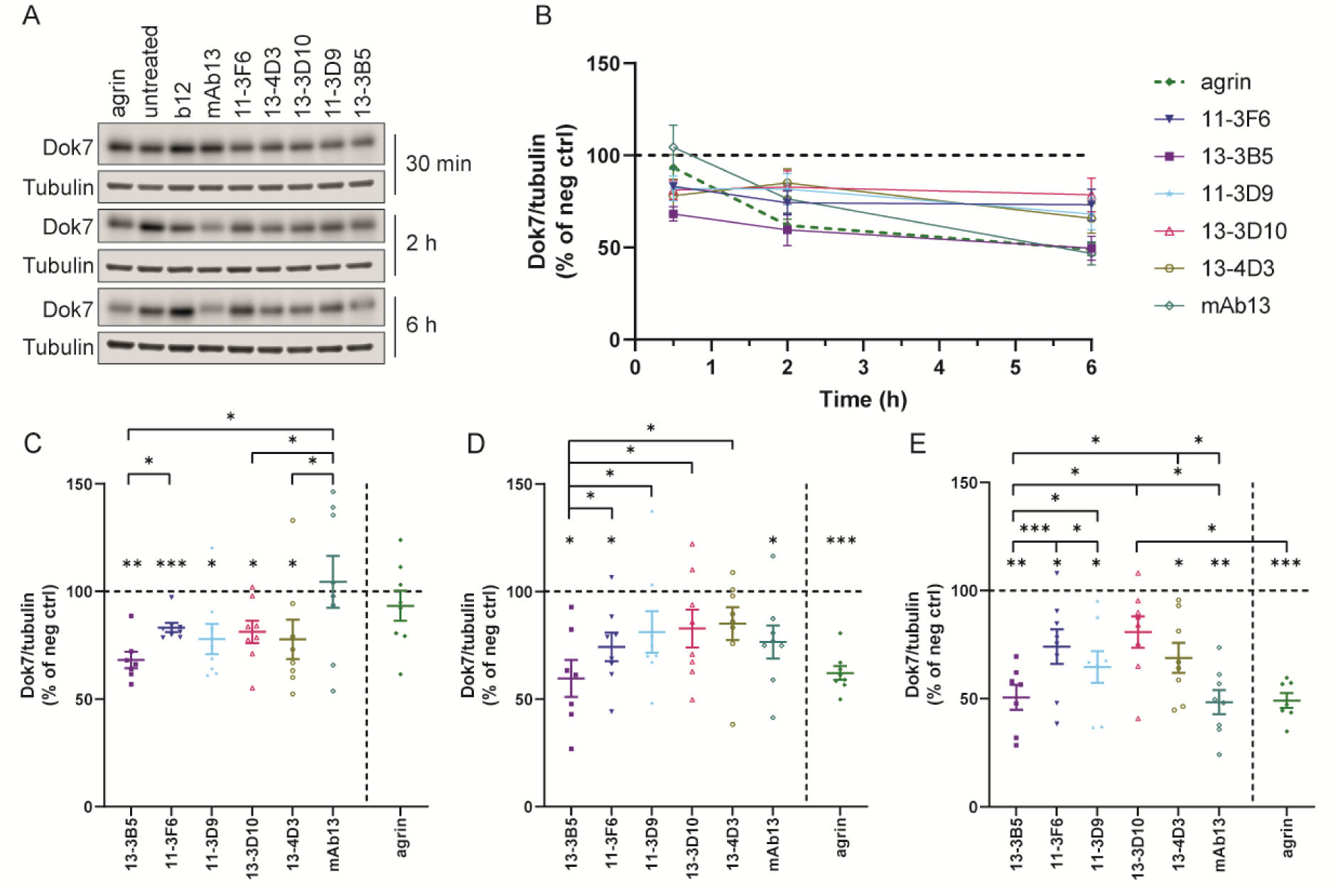
Bivalent MuSK antibodies affect Dok7 levels differently over time depending on epitope and clone. (A) Representative blots of Dok7 levels at 30 min, 2 h and 6 h. (B) Quantification of Dok7 levels over time, (C) after 30 min, (D) 2 h and (E) 6h. Data represents mean and SEM and is normalized against b12 for MuSK mAbs, and against untreated for agrin. n=8 (except for 13-3B5 n=7) for 30 min and 2 h, n=8 (except for untreated, agrin and 13-3B5 n=7) for 6h. Paired t-tests on log-transformed data with Benjamini-Hochberg false discovery rate correction. ∗p < 0.05; ∗∗p < 0.01; ∗∗∗p < 0.001.

### 3.4. MuSK antibodies do not deplete MuSK from the membrane of C2C12 myotubes

To gain a more detailed understanding of the factors that influence pathogenicity of MuSK antibodies, we studied the 13-3B5 and 11-3F6 clones further. These clones were selected since they differ in pathogenic capacity in bivalent form upon passive transfer to mice while binding the same domain of MuSK, preventing confounding effects of different target domains on MuSK [19]. Accelerated internalization is a common consequence of antigen-crosslinking by bivalent antibody binding. In addition, multiple receptor tyrosine kinases are known to rapidly internalize upon activation [31]. We hypothesized that if MuSK is depleted from the membrane by antibody binding, this can contribute to the pathogenicity of MuSK antibodies. To investigate whether the amount of endogenously expressed MuSK on the membrane surface is affected by exposure to MuSK antibodies, we biotinylated and pulled down membrane surface proteins of C2C12 myotubes and probed this fraction for MuSK immunoreactivity. Neither monovalent nor bivalent MuSK antibodies binding the Ig-like domain 1 reduced the amount of cell surface MuSK after 30 minutes of exposure (Figure 4). Agrin, at the minimal dose for maximal activation (0.11 nM) or a supramaximal dose (5 nM), also did not reduce the amount of cell surface MuSK. Exposure to epidermal growth factor (EGF) for 15 minutes did significantly reduce the amount of cell surface EGF receptor in C2C12s, validating the method. Longer exposure to agrin or bivalent 13-3B5 also did not reduce endogenous MuSK on the surface membrane (Appendix A, Figure S4). Thus, the amount of MuSK on the membrane seems stable during active signaling and is not altered by exposure to these patient-derived MuSK antibodies in C2C12 myotubes.

**Figure 4:**
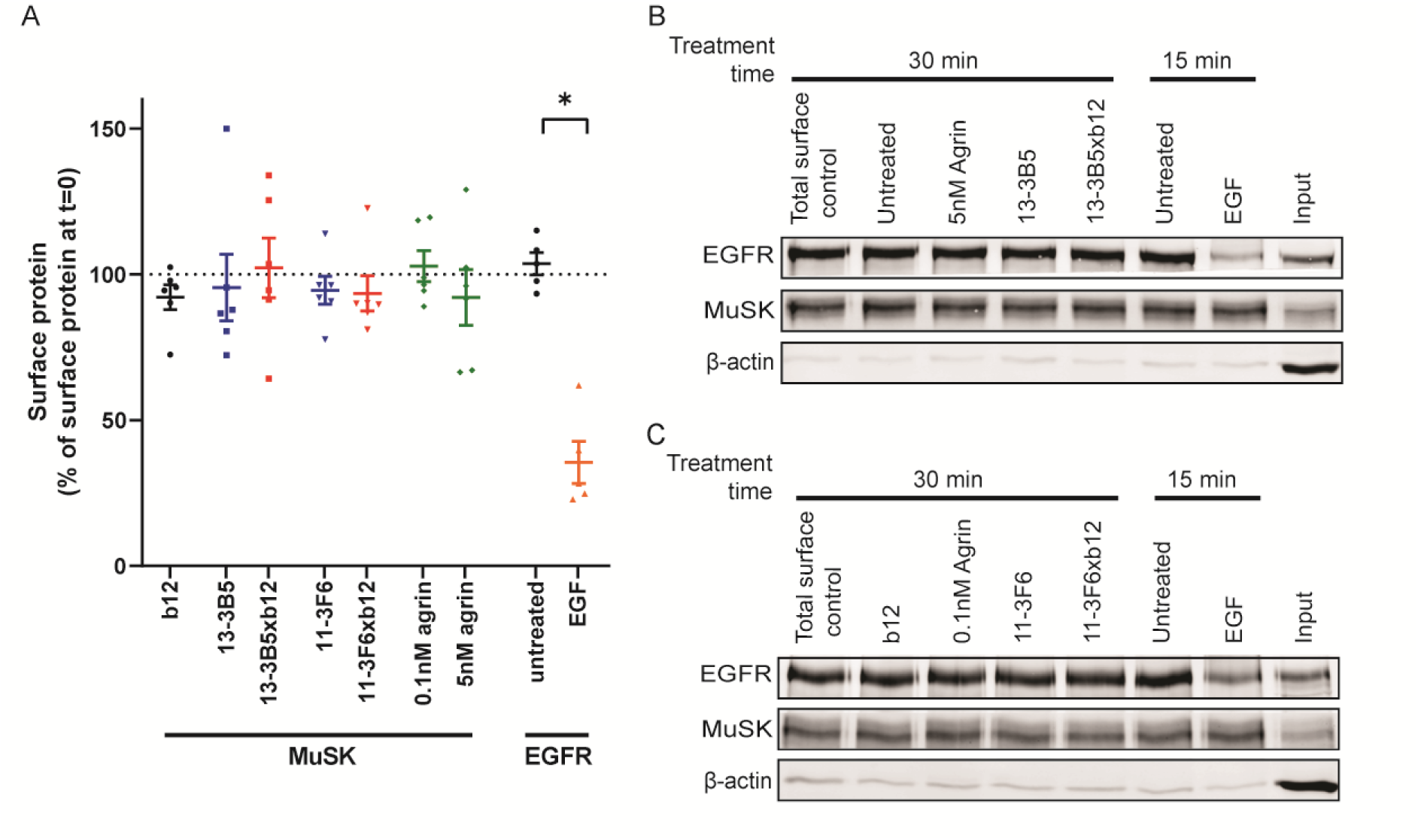
MuSK surface depletion does not occur following exposure to monovalent or bivalent MuSK antibodies. (A) Quantification of surface MuSK following 30 min exposure to monovalent or bivalent MuSK antibodies or agrin compared to t=0 in C2C12 myotubes (n=6). Significant surface depletion of the EGFR following EGF exposure could be detected with this method (n=5). Representative blots of surface MuSK and EGFR upon (B) 13-3B5(xb12) or (C) 11-3F6(xb12). Intracellular protein β-actin is not pulled down with this method. Data depicts mean ± SEM. Paired t-test on log-transformed data with Holm-Bonferroni correction. ∗p < 0.05.

### 3.5. Neuromuscular junction gene expression is differentially affected by monovalent and bivalent MuSK antibodies

Expression of synaptic genes for the specialized structure and function of the NMJ is tightly regulated and specific to subsynaptic nuclei of muscles [7]. Disruptions in their expression may contribute to the pathomechanism of disease-causing MuSK antibodies. To investigate if altered NMJ gene expression may explain the pathogenic differences observed between MuSK antibodies, we measured RNA expression of NMJ genes with a direct link to MuSK signaling in the masseter muscle of NOD/SCID mice exposed for 11 days or 3 weeks to bivalent or monovalent MuSK antibodies [19]. The masseter is a bulbar muscle, which displays prominent weakness in MuSK MG patients [32]. Briefly, monovalent MuSK antibodies to the Ig-like domain 1 caused severe myasthenic muscle weakness, lethal after 11 days (Figure 5A). In contrast, bivalent 11-3F6 did not cause overt muscle weakness after 3 weeks, while exposure to bivalent 13-3B5 resulted in subclinical myasthenic muscle weakness after 11 days progressing to lethal muscle weakness after 3 weeks [19].

**Figure 5:**
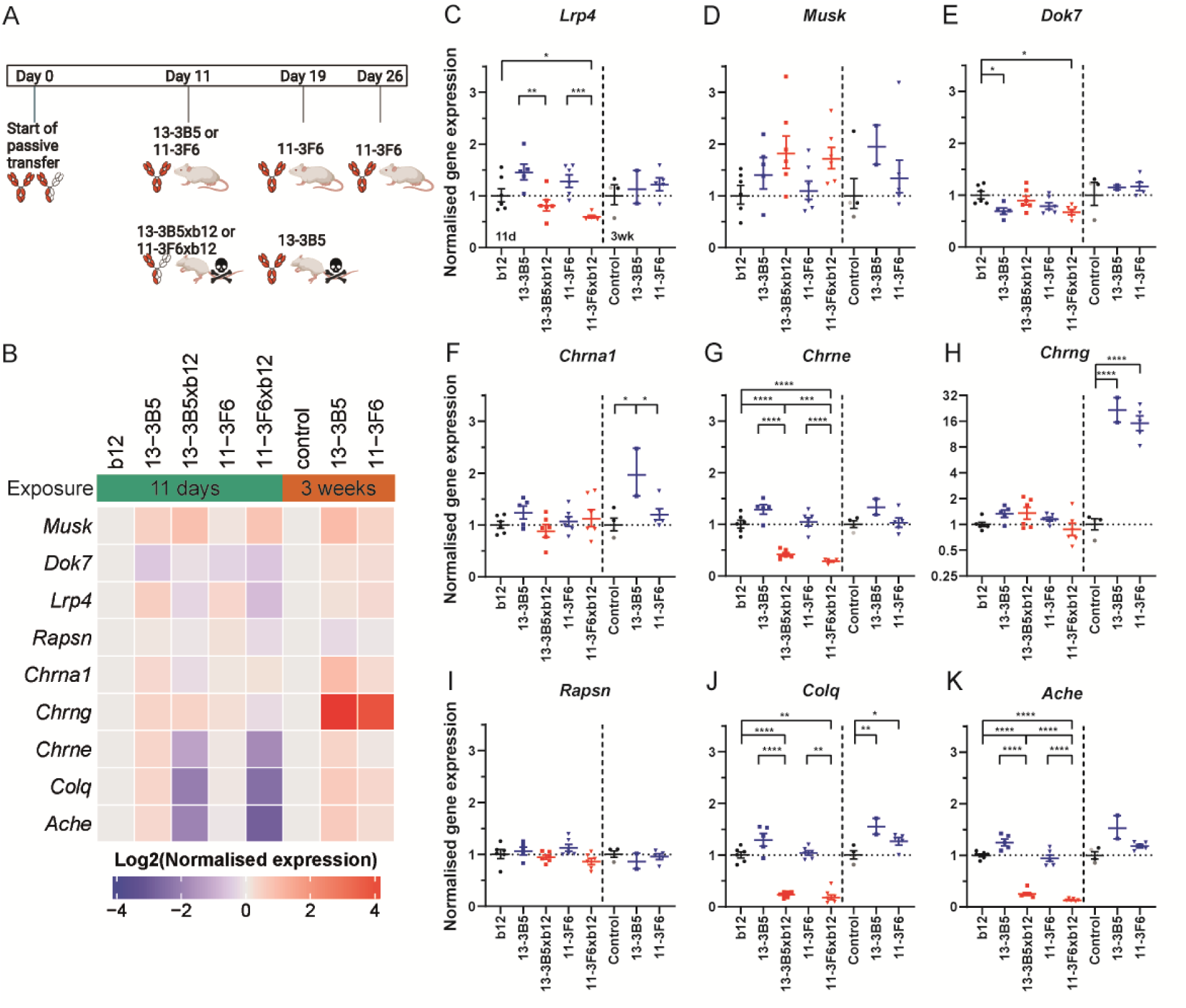
Monovalent and bivalent MuSK antibodies have differential effects on NMJ gene expression in mouse muscle. (A) Graphic summary of results and time frame of passive transfer experiments ^16^. (B) Log2 normalized expression of NMJ genes in masseter muscle of NOD/SCID mice exposed to MuSK antibodies. Normalized gene expression of (C) Lrp4, (D) Musk, (E) Dok7, (F) Chrna1, (G) Chrne, (H) Chrng, (I) Rapsn, (J) Colq, (K) Ache in masseter muscle of exposure to MuSK antibodies for 11 days (left of dotted line) or 3 weeks (right of dotted line). Data depicts geometric mean ± geometric SEM. 11 days: 2.5mg/kg b12, 13-3B5xb12, 11-3F6, 11-3F6xb12: n=6; 2.5mg/kg 13-3B5: n=5. 3 weeks: control is 10mg/kg b12 (black): n=2 combined with untreated or PBS-treated (grey): n=2; 10mg/kg 13-3B5: n=2; 10mg/kg 11-3F6: n=5. One-way ANOVA with Šidák-corrected comparisons for 11 day exposure. One-way ANOVA with Fisher’s LSD test for 3 week exposure. Welch ANOVA when the assumption of equal variance was not met. *p < 0.05, **p < 0.01, ***p < 0.001, ****p < 0.0001

Monovalent and bivalent MuSK antibodies caused different patterns of NMJ gene expression in the masseter muscle. In general, monovalent MuSK antibodies showed a (tendency to) decreased expression of a subset NMJ genes, while bivalent MuSK antibodies showed (a tendency to) increased expression of a partially overlapping subset of NMJ genes (Figure 5B). MuSK antibodies differentially affected the expression of *Lrp4*, *Chrne*, *Chrng*, *Colq* and *Ache* depending on valency. Most notably, monovalent MuSK antibodies strongly downregulated *Colq*, *Ache* and the epsilon subunit of the AChR (*Chrne*), while bivalent MuSK antibodies even after prolonged exposure did not (Figure 5G, J and K). In contrast, both bivalent MuSK antibodies strongly increased the expression of the gamma subunit of the AChR (*Chrng*), while this was not caused by exposure to monovalent MuSK antibodies at end-stage disease (Figure 5H). The alpha 1 subunit of the AChR (*Chrna1*) was only increased by bivalent 13-3B5 after 3 weeks of exposure (Figure 5F). Lastly, *Musk* is the only gene with a trend (p=0.07) to increased expression for all MuSK antibodies (both monovalent and bivalent) that caused a myasthenic phenotype (Figure 5D). Taken together, these differential effects on NMJ-specific gene expression further support monovalent and bivalent MuSK antibodies cause myasthenic muscle weakness through different mechanisms.

To investigate if MuSK antibody binding directly affects (NMJ) gene expression in muscle, we performed a transcriptome analysis of C2C12 myotubes exposed to agrin, bivalent 13-3B5, or agrin in combination with monovalent 13-3B5xb12 or 11-3F6xb12 for 16h, as at this time the full AChR clustering cascade is active. No differentially expressed genes were found upon MuSK activation or MuSK-antibody binding (see Appendix B and C). To confirm we did not miss the gene expression window induced by MuSK signaling, we exposed C2C12s to agrin or bivalent MuSK antibodies for different durations between 30 minutes and 24 hours and measured the expression of *Chrne* and *Musk*, because their promotors have experimentally been shown to be directly regulated by MuSK signaling [33, 34]. MuSK activation by agrin or bivalent MuSK antibodies did not affect *Chrne* or *Musk* expression within 24 hours (Appendix A, Figure S5A and B). Neuregulin activates the other pathway regulating NMJ gene expression in muscles and did rapidly increase the expression of early growth response 3 (*Egr3*) and showed a tendency to increase *Chrne* expression after 24 hours (Appendix A, Figure S5B and C) [7]. This suggests gene expression changes relevant for the NMJ can be detected in C2C12 myotubes, however, those mediated through MuSK may require more time. Exposure to MuSK activating compounds for more than 24 hours resulted in cell death, hindering the investigation of longer-term or indirect effects of MuSK signaling on gene expression. Taken together, the observed changes in NMJ gene expression in the clinically relevant masseter muscle are unlikely to be mediated by an acute effect of MuSK-antibody binding on MuSK-mediated modulation of gene expression.

## 4. Discussion

To further understand the mechanisms underlying MuSK MG, we tested how a panel of monovalent and bivalent (patient-derived) monoclonal MuSK antibodies affected MuSK-mediated signaling. The valency and/or epitope of these MuSK antibodies had a significant effect on antibody binding kinetics, MuSK activation, Dok7 levels and NMJ gene expression. Together, these data support that pathogenic monovalent and bivalent MuSK antibodies cause NMJ dysfunction through different molecular processes that depend on epitope, highlighting the heterogeneity and complexity of the disease mechanisms underlying MuSK MG (Figure 6).

**Figure 6:**
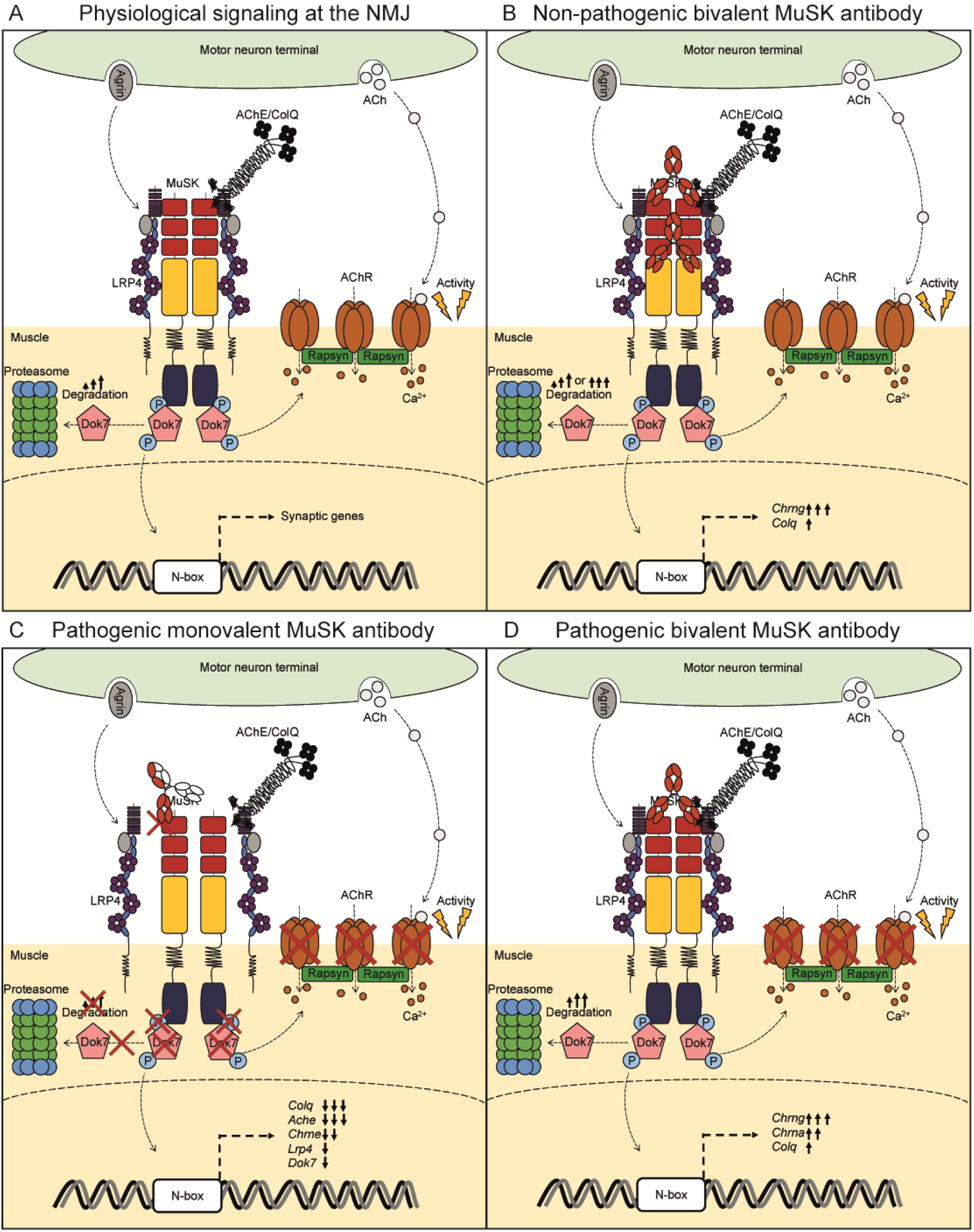
Model of how MuSK antibodies with different functional characteristics impact signaling at the NMJ. (A) During physiological signaling agrin-Lrp4 dimerizes and autophosphorylates MuSK. Dok7 binds to phosphorylated MuSK and is phosphorylated itself initiating a signaling cascade leading to clustering of AChR. In addition, MuSK binds ColQ-AChE complex, tethering it to the synapse, and MuSK is involved in the expression of synaptic genes. Activity-dependent degradation of Dok7 over time may be the break of this signaling cascade. (B) Non-pathogenic bivalent MuSK antibodies have some effects on MuSK signaling, but these are not sufficient to cause dysfunction of neurotransmission. Pathogenic (C) monovalent and (D) bivalent MuSK antibodies have completely different effects on MuSK functioning, though both ultimately cause declustering of AChRs and myasthenic symptoms.

Monovalent MuSK antibodies cause NMJ dysfunction by inhibiting MuSK-mediated signaling (Figure 6C). The monovalent MuSK antibodies binding the Ig-like domain 1 bound strongly to MuSK, inhibited agrin-induced MuSK phosphorylation and Dok7 binding to MuSK. Previously, monovalent Fabs of monoclonal MuSK antibodies binding the Ig-like domain 2 fully inhibited agrin-induced AChR clustering, similar to these Ig-like domain 1 binders [18, 19, 35, 36]. Polyclonal patient IgG4 also blocks agrin-induced MuSK activation and AChR clustering, mainly targets the N-terminal Ig-like domains 1 and 2 of MuSK, and is estimated to be up to 99% monovalent due to Fab-arm exchange [12, 17, 21, 37]. In line with these *in vitro* effects, polyclonal patient IgG(4) and monovalent MuSK monoclonal antibodies targeting the MuSK Ig-like domain 1 cause myasthenia in mice [19, 38, 39]. Monovalent mAb13xb12 did not inhibit MuSK activation nor Dok7 binding compared to agrin, suggesting monovalent binding to the Fz domain may not be pathogenic *in vivo*. Taken together, inhibiting the MuSK-mediated AChR clustering pathway is an important mechanism of disease for monovalent MuSK antibodies and this may depend on the MuSK domain targeted.

Monovalent MuSK antibodies also have consequences beyond the canonical MuSK-mediated AChR clustering pathway. Further downstream, monovalent MuSK antibodies binding the Ig-like domain 1 also decreased the gene expression of *Colq*, *Ache* and *Chrne* and to a lesser extent *Lrp4* in mouse muscle. This suggests a production shortage of both mature AChRs and the ColQ-AChE complex that may impair proper functioning. Disruption of the ColQ-AChE complex may contribute to the deleterious effects of AChE inhibitors in MuSK MG patients and passive transfer models [40, 41]. Additionally, ColQ binds to Lrp4 and antibodies blocking MuSK-Lrp4 interaction may impair ColQ tethering to the postsynaptic structure [8]. The inhibition of MuSK by monovalent antibodies thus has consequences throughout the NMJ that may contribute to clinical symptoms and treatment responsiveness.

In contrast to monovalent MuSK antibodies, bivalent MuSK antibodies are able to induce the early steps of MuSK signaling independent of agrin. Bivalent MuSK antibodies against the Ig-like domain 1 or the Fz domain induced MuSK phosphorylation and Dok7-binding to MuSK. This is in line with previous observations that monoclonal and polyclonal anti-MuSK IgG can induce MuSK, Dok7 and/or AChRβ phosphorylation similar to agrin, independent of the MuSK domain targeted [17-20, 23, 35, 42]. In sum, binding site of agonist MuSK antibodies does not affect their ability to force dimerization and initiate the early steps of MuSK-mediated signaling.

Cellular Dok7 levels do appear to be differentially regulated depending on how MuSK is activated. Bivalent MuSK antibodies to the Ig-like domain 1 reduce Dok7 levels faster compared to bivalent mAb13 (binding the Fz domain) and agrin. In addition, bivalent 13-3B5 resulted in the largest reduction of Dok7 compared to the other Ig-like domain 1 binding clones at all time points. Dok7 is essential for forming and maintaining the NMJ and reduction of Dok7 levels by the proteosome is thought to be a crucial negative feedback loop in regulating agrin-induced MuSK signaling [5, 20, 43, 44]. Bivalent 13-3B5 has the strongest binding to MuSK, has a non-overlapping epitope compared to the other Ig-like domain 1 binders, and causes muscle weakness *in vivo* [19, 24]. The early and relatively strong reduction of Dok7 levels by bivalent 13-3B5 may indicate Dok7 is degraded too much too quickly upon activation, hampering its function (Figure 6D). This was also suggested in earlier experiments using polyclonal bivalent anti-MuSK IgG from rabbits [20]. Together, these results suggest that the kinetics of Dok7 degradation may depend on how MuSK is dimerized depending on affinity and/or antibody-epitope between and within structural MuSK domains. However, since agrin induces the same reduction of Dok7 levels after 6 hours as bivalent 13-3B5, the kinetics and consequences for downstream signaling should be investigated further to understand if rapid Dok7 depletion contributes to the *in vivo* pathogenicity of bivalent 13-3B5.

Bivalent MuSK antibodies also showed a completely different signature on NMJ gene expression in mouse masseter muscle compared to monovalent MuSK antibodies. Bivalent MuSK antibodies increased the expression of *Chrng, Chrna1, Colq,* and to a lesser extent *Musk*, *Ache* and *Chrne* after three weeks, similar to what was seen upon active immunization with MuSK [45]. Active immunization with MuSK induces bivalent MuSK antibodies, because murine antibody subclasses cannot undergo Fab-arm exchange *in vivo.* Active immunization models of MuSK MG thus only recapitulate the disease mechanisms mediated by bivalent MuSK antibodies. Presynaptic denervation is associated with a strong upregulation of *Chrng* expression and has been observed at end-stage disease upon active immunization with MuSK and passive immunization with patient-derived IgG(4) [38, 39, 46, 47]. However, abnormalities in presynaptic morphology due to bivalent 13-3B5 (pathogenic) were not observed at symptom onset at 11 days (congruent with the lack of *Chrng* expression at that time), while postsynaptic pathology was already present in both a bulbar and a limb muscle [19]. In addition, *Chrng* expression was also upregulated by bivalent 11-3F6, which does not cause muscle weakness even after months of exposure [24]. Taken together, if the upregulation of *Chrng* expression by bivalent MuSK antibodies after three weeks is a sign of presynaptic pathology, it is unlikely to fully drive the pathogenicity of bivalent MuSK antibodies.

Finally, rapid antibody-mediated or activity-dependent surface depletion of MuSK is also unlikely to be a major part of the pathomechanism of bivalent MuSK antibodies. Endogenous MuSK did not deplete from the surface membrane upon treatment with bivalent or monovalent MuSK or agrin in C2C12 cell cultures. MuSK internalization from the cell surface has not been consistently found upon agrin or MuSK antibody treatments [21, 31, 48-50]. This may be due to experimental differences, such as 1) agrin and/or MuSK antibodies with different properties, 2) cell culture systems endogenously or exogenously expressing MuSK, 3) how MuSK is measured or 4) whether new protein synthesis is inhibited. Gemza et al. furthermore elegantly discuss the potential slow rate of MuSK endocytosis in relation to previous studies and other receptor tyrosine kinases may be due to 1) the indirect activation of MuSK through Lrp4 and Dok7 and 2) anchoring of MuSK in the dense cytoskeletal structure of the post-synapse [31, 48, 49]. In sum, a clear mechanism of how bivalent MuSK antibodies disrupt the organized structure of the NMJ remains elusive. The extracellular domain of MuSK is also important for AChR clustering through an undiscovered process independent of the kinase domain of MuSK [51]. This unidentified function may play an important role in the pathomechanisms of bivalent MuSK antibodies.

Taken together, MuSK MG patients have a polyclonal antibody response resulting in a mixture of MuSK antibodies against different epitopes, and of IgG subclasses differing in functional valency and complement activating capacity [10, 11, 14]. By dissecting the role of antibody epitope, affinity, and signalling effects, this study contributes to an increased understanding on how these variables affect the pathophysiology in MuSK MG patients. The net effect of MuSK autoantibodies in individual patients will be determined by the unique composition of MuSK antibodies and their characteristics, in combination with their relative titres. It will be interesting to get a more in dept view on the heterogeneity and clonality of the MuSK autoantibody response in individual patients and to study if these characteristics can explain the inter-individual discrepancy between autoantibody titer and severity of the clinical symptoms [52, 53]. Additionally, several non-pathogenic, agonistic MuSK antibodies have shown therapeutic benefits in pre-clinical models of several neuromuscular diseases with impaired neuromuscular junction integrity [15, 23-25, 30, 54-56]. This study sheds light on how bivalent MuSK antibodies with different characteristics impact MuSK-mediated signaling further aiding therapeutic development of agonistic MuSK antibodies.

## Supporting information

Appendix B Differential Expression

Appendix C Counts

## Acknowledgements

We thank Anita van den Heuvel for help with the RNAseq analyses, Laurent Paardekooper, Oscar Dekker, Jessica van Bokkum and Manon van den Bout for help with designing and producing recombinant MuSK and William Jiemy for constructive feedback on the manuscript. The LUMC forms part of the European Reference Network for Rare Neuromuscular Diseases (ERN EURO-NMD) and the Netherlands Neuromuscular Disorders Center (NL-NMD).

