## Appendix A: Supplementary Materials for "MuSK antibodies differently affect the MuSK signaling cascade depending on valency and epitope specificity"

### Supplementary information

A

| Clone | 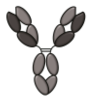 | 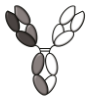 | 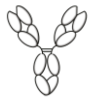 |
|---|---|---|---|
| 13-4D3 | 2.1% | 82.2% | 15.7% |
| 13-3D10 | 1.1% | 97.1% | 1.8% |
| mAb13 | 1.7% | 97.0% | 1.1% |

B

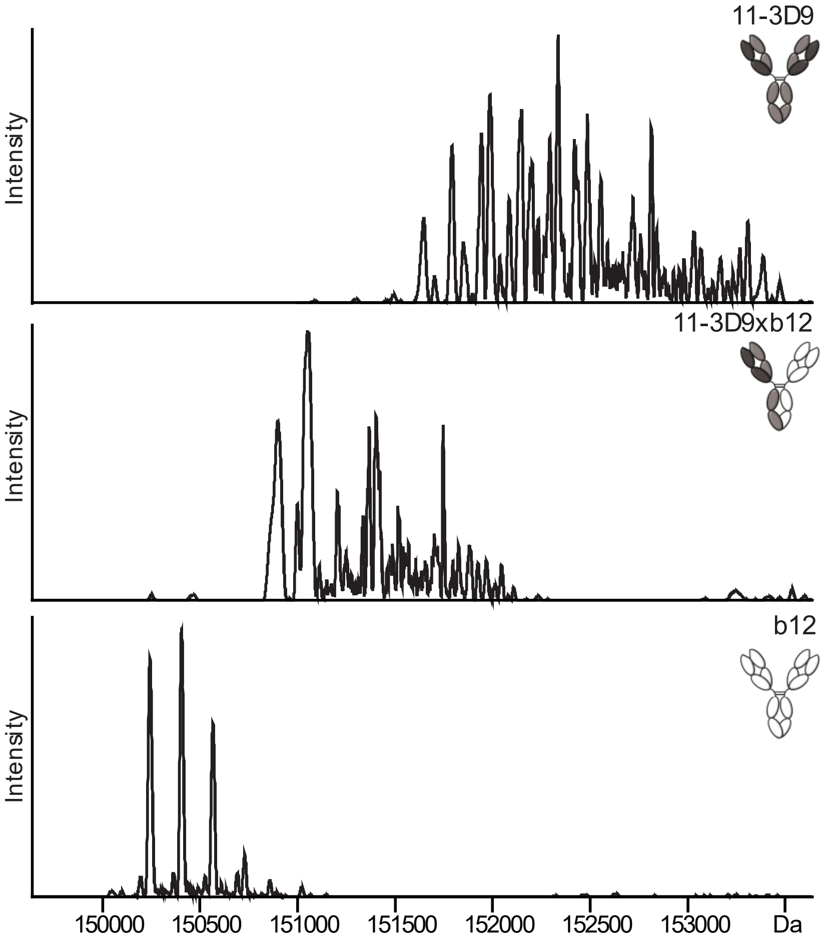


*Figure S1: Exchange efficiency of monovalent MuSK antibodies. (A) Relative amounts of bivalent monospecific and monovalent bispecific antibody variants after the controlled Fab-arm exchange (cFAE) reaction measured by capillary electrophoresis. (B) Deconvoluted mass spectra show generation of monovalent bispecific 11-3D9xb12 after cFAE.*

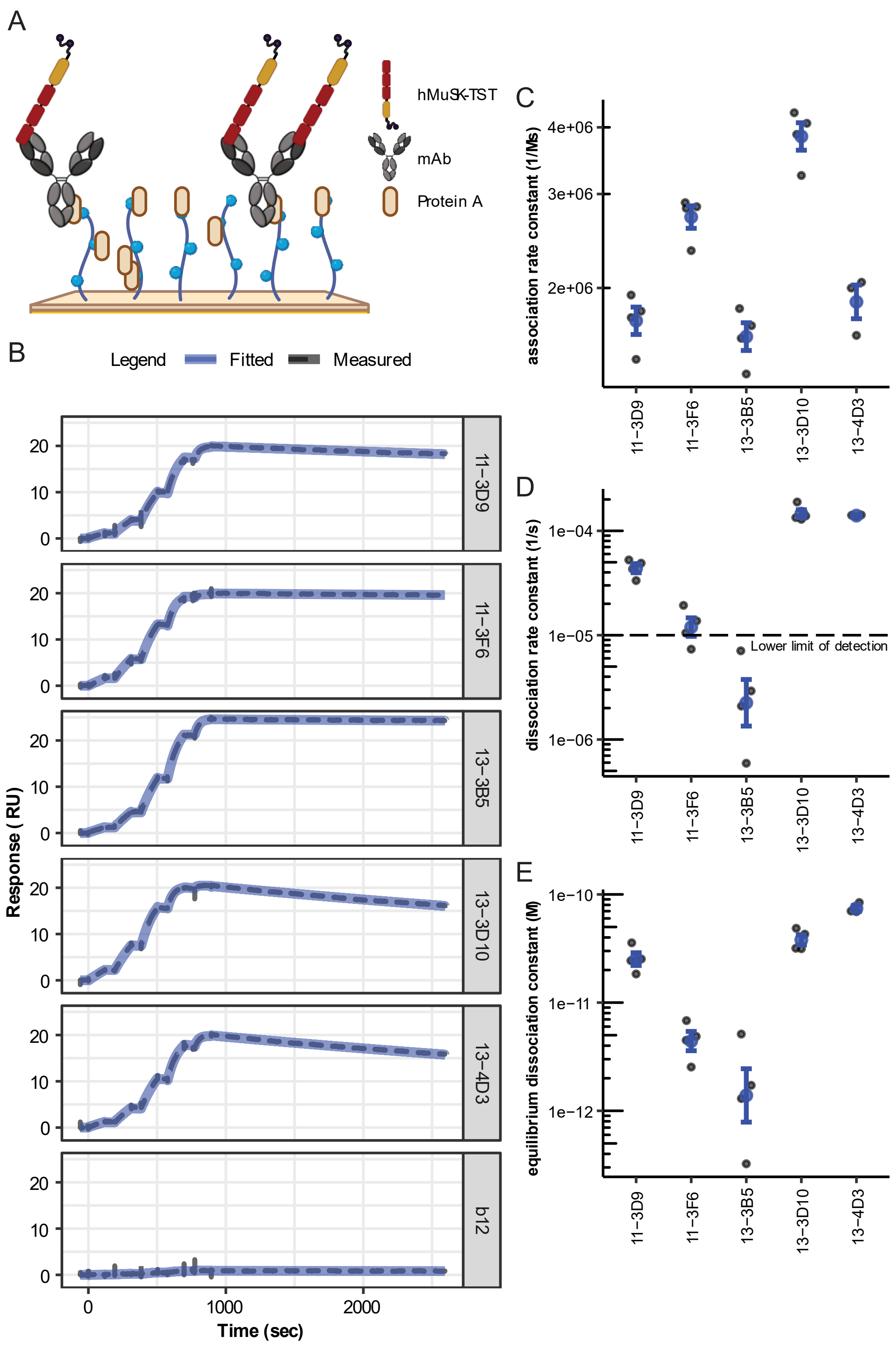


*Figure S2: Binding kinetics of MuSK to MuSK antibodies. (A) Graphical representation of SPR assay with mAbs as a ligand, captured by protein A, and hMuSK-TST as analyte. (B) Representative double referenced sensorgram with the measured response (black dashed) and 1:1 fitted response (solid blue) of MuSK binding to MuSK antibodies. (C) Association rate constant, (D) dissociation rate constant and (E) equilibration dissociation constant fitted with a 1:1 model. Data represents geometric mean and geometric SEM of n=4 independent replicates (n=3 for 13-4D3).*

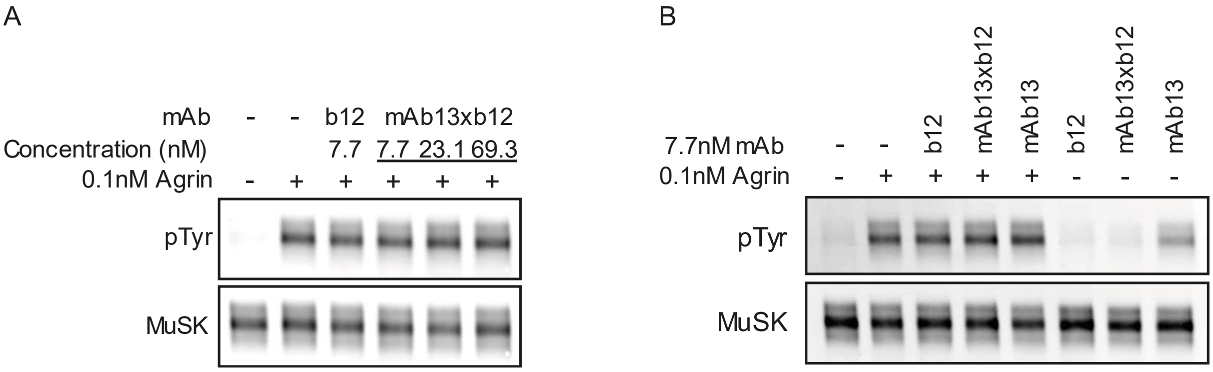


*Figure S3: Monovalent mAb13xb12 does not inhibit agrin-induced MuSK phosphorylation. (A) Agrin-induced MuSK phosphorylation in combination with increasing concentrations of mAb13xb12. (B) MuSK phosphorylation upon addition of monovalent mAb13xb12 or bivalent mAb13 in the absence or presence of agrin.*

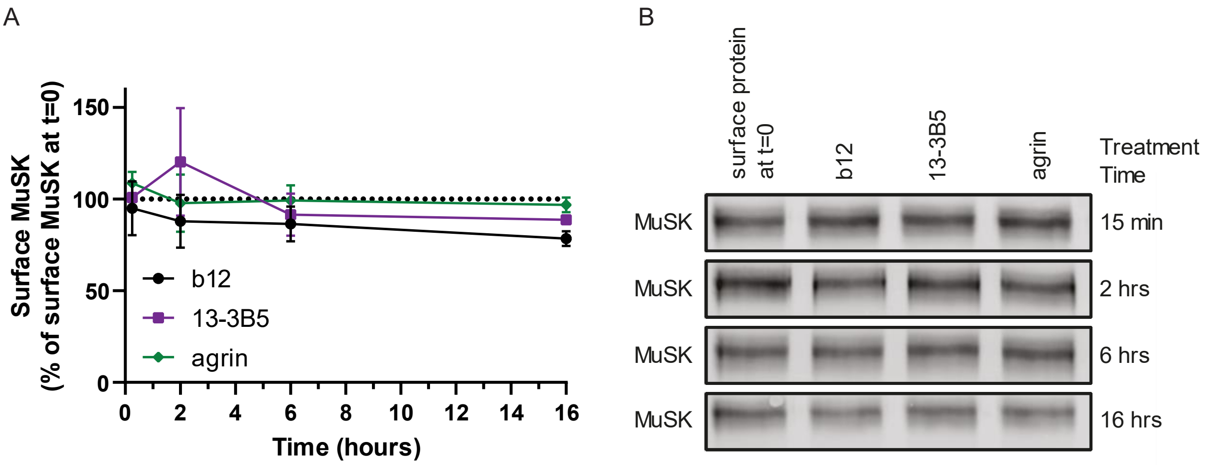


*Figure S4: Longer exposure to agrin or bivalent 13-3B5 does not reduce surface MuSK. (A) Surface MuSK does no significantly differ over time upon exposure to agrin or bivalent 13-3B5. (B) Representative blot of surface MuSK after 15 min, 2 h, 6 h or 16 h exposure. Data represent mean ± SEM over n=2 (15min, 2h and 6h) or n=4 (16h) experiments.*

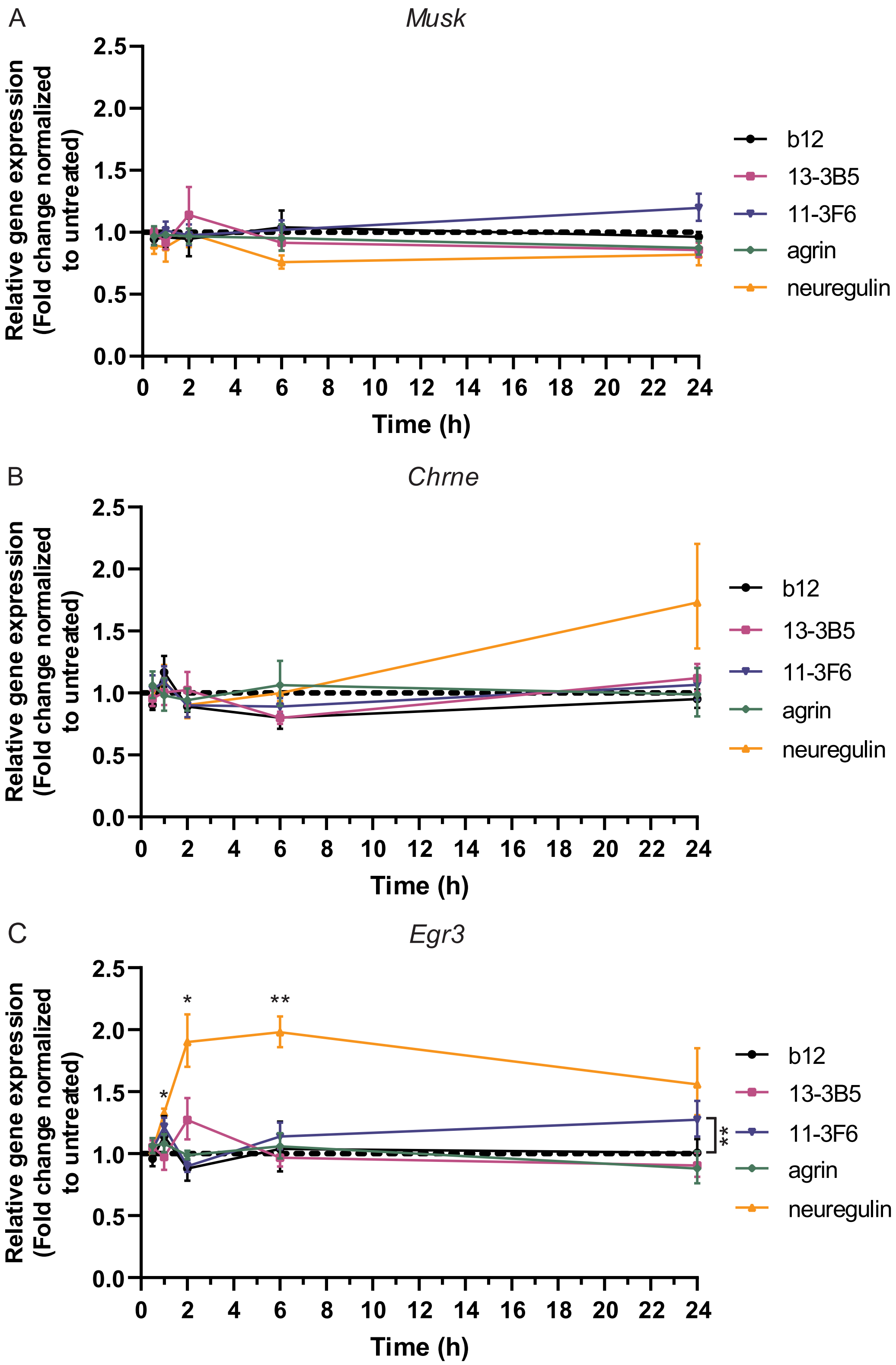


*Figure S5: MuSK signalling does not directly induce gene expression of Musk or Chrne in C2C12 myotubes. Normalised gene expression of (A) Musk, (B) Chrne and (C) Egr3. Egr3 expression is regulated by neuregulin and serves as a positive control for the method. 30 min (n=5), 60 min (n=4), 2 h (n=5), 6 h (n=5) and 24 h (n=6). Data depicts geometric mean ± geometric SEM. Paired t-test on log2-transformed data with Benjamini-Hochberg false discovery rate correction. *p < 0.05, **p < 0.01.*

*Table S1: Antibody conditions for western blot*

| Antigen | Block | Antibody buffer | Primary antibody | Secondary antibody |
|---|---|---|---|---|
| MuSK IP samples | | | | |
| MuSK | 3% BSA | Immunobooster | AF562<br>R&D systems<br>0.2µg/mL | 926-32214<br>Licor<br>0.2 µg/mL |
| Phospho-MuSK | 3% BSA | Immunobooster | 05-321<br>Millipore<br>1µg/mL | 926-68072<br>Licor<br>0.2 µg/mL |
| Dok7 | 3% BSA | 0.5% BSA | AF6398<br>R&D systems<br>0.5µg/mL | 205-032-176<br>Jackson ImmunoResearch<br>1:10.000 |
| Whole cell lysate | | | | |
| Dok7 | 5% milk | 2% milk | AF6398<br>R&D systems<br>0.5µg/mL | 205-032-176<br>Jackson ImmunoResearch<br>1:10.000 |
| Tubulin | 5% milk | 2% milk | T6199<br>Sigma-Aldrich<br>0.2µg/mL | 926-68072<br>Licor<br>0.2 µg/mL |
| Surface depletion assay | | | | |
| MuSK | Odyssey Blocking Buffer | Immunobooster | AF562<br>R&D systems<br>0.2µg/mL | 926-32214<br>Licor<br>0.1 µg/mL |
| EGFR | Odyssey Blocking Buffer | Immunobooster | 51091-T52<br>Bioconnect<br>1:1000 | 926-32213<br>Licor<br>0.1 µg/mL |
| β-actin | Odyssey Blocking Buffer | Immunobooster | ab8226<br>Abcam<br>0.2µg/mL | 926-68072<br>Licor<br>0.1 µg/mL |

*Table S2: Primers with validated amplification efficiency in C2C12 myotubes and NOD/SCID masseter muscle*

| **Gene** | **Accession number** | **Primer sequence** | **Amplification size (bp)** | **Amplification efficiency (%) C2C12** | **Correlation coefficient ($R^2$) C2C12** | **Amplification efficiency (%) Masseter** | **Correlation coefficient ($R^2$) Masseter** |
|---|---|---|---|---|---|---|---|
| *Gapdh* | MGI:95640 | Fw: TCCATGACAACTTTGGCATTG<br>Rv: TCACGCCACAGCTTTCCA | 103 | 104,3 | 0,998 | 108,8 | 0,996 |
| *Rpl13a* | MGI:1351455 | Fw: TGCTGCTCTCAAGGTTGTTC<br>Rv: TTCTCCTCCAGAGTGGCTGT | 114 | 98,1 | 1,000 | 104,9 | 0,994 |
| *Egr3* | MGI:1306780 | Fw: CTGACAATCTGTACCCCGAGGA<br>Rv: GCTTCTCGTTGGTCAGACCGAT | 129 | 105,2 | 0,997 | | |
| *Musk* | MGI:103581 | Fw: AACCCCAAACCATCTGTGTC<br>Rv: GTCCTGCATCTTCCTTTTGC | 121 | 101,1 | 0,991 | 104,6 | 0,991 |
| *Chrne* | MGI:87894 | Fw: GAACTCGTGTTTGAGGGTCAG<br>Rv: TCAGCCACAAAGTTCACAGC | 125 | 103,8 | 0,993 | 96,9 | 0,994 |
| *Chrna1* | MGI:87885 | Fw: AAGCACCCTGAGGTGAAAAG<br>Rv: CCATCACCATGGCAACATAC | 118 | | | 96,6 | 0,992 |
| *Chrng* | MGI:87895 | Fw: ATCGTCGTGAACTCTGTGGT<br>Rv: CCTTCCTCTCGAGCCATGAT | 215 | | | 98,1 | 0,983 |
| *Lrp4* | MGI:2442252 | Fw: ATGGGTCTATGCGGAAAGTG<br>Rv: CGCTCTAATTTGGCGTTCTC | 121 | | | 103,4 | 0,991 |
| *Dok7* | MGI:3584043 | Fw: TCAGCCTCAGAAGAGCGTGTTG<br>Rv: GCCTCAGAAGAGGAACTGGATAG | 137 | | | 108,7 | 0,986 |
| *Colq* | MGI:1338761 | Fw: TGTGGTCAACAACCAGGAAG<br>Rv: AAAGATCGCTGGTCTCTTCG | 78 | | | 100,4 | 0,991 |
| *Ache* | MGI:87876 | Fw: GGGCTCCTACTTTCTGGTTTACG<br>Rv: GGGCCCGGCTGATGAG | 71 | | | 102,4 | 0,997 |
| *Rapsn* | MGI:99422 | Fw: AGGCTGGAGCCTCAAATATC<br>Rv: AGGGCAATCTTCATGGACTC | 117 | | | 107,4 | 0,997 |

*Table S3: p-values of Tukey multiple comparisons of clones on the apparent $k_d$*

| **Clone** | **11-3F6** | **11-3D9** | **13-3D10** | **13-4D3** |
|---|---|---|---|---|
| **13-3B5** | <0.0001 | <0.0001 | <0.0001 | <0.0001 |
| **11-3F6** | | <0.001 | <0.001 | <0.01 |
| **11-3D9** | | | 0.947 | 0.968 |
| **13-3D10** | | | | 0.657 |
